# Research advancements on the application of sterile insect technique for the control of lepidopteran pests over the past two decades: A Systematic Review

**DOI:** 10.1101/2025.01.17.633492

**Authors:** Bambo Karabo, LN Malinga

**Affiliations:** South African Sugarcane Research Institute, Entomology Department, Mount Edgecombe, Durban, KwaZulu-Natal, South Africa; University of KwaZulu-Natal, School of Life Sciences, Scottsville, Pietermaritzburg, KwaZulu-Natal, South Africa

**Keywords:** SIT, Moths, Gamma, X-ray, Lepidoptera

## Abstract

Many lepidopterans’ species, such as fruit borer (*Helicoverpa armigera*) on tomato, apple moth *Cydia pomonella* (codling moth), and orange moth *Thaumatotibia leucotreta* (false codling moth), are among many lepidopterans that are crop pests. In the early 1930s, the concept of the sterile insect technique was conceived in response to the global threat of insect pests and the excessive use of insecticides, which were proven to pose a health risk to humans. This pest control tactic has gained and is currently gaining more traction in all regions where it is implemented on both small and large scales. This systematic review looks at the recent research advances in the application of sterile insect techniques for the control of lepidopteran pests. The paper analyses publications, focusing on the geographic distribution of studies, target species, irradiation source, and the year of publication. It highlights the growing interest in SIT as an environmentally friendly pest management strategy compared to insecticide use. The paper compiles and summarises information over the last two decades on the lepidopteran species studied, the study approach, the country where the study was performed, the source of irradiation, and the year of publication. It also analysed the suitability of each lepidopteran species as an SIT candidate and the effectiveness of this control strategy. The review resulted in 2 537, from which 24 publications matched the search criteria. From the results, 14 species were identified, and the most studied lepidopteran was *T. leucotreta*. The review discusses the effectiveness of SIT in controlling various lepidopteran species and explores the potential for further development and implementation of SIT programmes globally.

## INTRODUCTION

Most lepidopteran species are responsible for causing serious damage to agricultural crops worldwide (Ouaba *et al*., 2022). They can be pests of great economic value and are among the most damaging to food and fibre crops (José *et al*., 2022). Currently, these pests are controlled using a broad range of insecticides that are persistently sprayed in large quantities (Liu *et al*., 2023). However, the use of insecticides for the control of pests is discouraged because it is unhealthy for the environment (Pérez-staples *et al*., 2021; Zhao *et al*., 2022), have negative effects on the health of humans, and huge quantities are needed to continuously spray the crops, making it economically unfavourable (Bourtzis and Vreysen, 2021; Vreysen *et al*., 2016). Lepidopterans often also rapidly develop resistance to every newly developed chemical insecticide, which also causes further challenges to sustainable control (Osouli *et al*., 2021). As more scientific evidence has shown the long-term effects of insecticides on human health and the environment, biological control tools such as the sterile insect technique (SIT) have gained more traction.

For the past six to seven decades, SIT has been part of the area-wide integrated pest management (AW-IPM) strategies to suppress insect pest populations (Bourtzis and Vreysen., 2021; Dyck *et al*., 2021). Area-wide integrated pest management can be defined as a systematic reduction of a target key pest to predetermined population levels through uniformly applied control measures over a large geographical area clearly defined by biologically based criteria (Klassen & Vreysen, 2020). The control tactics of this programme target the entire insect population and are integrated based on species specificity (Dalal *et al*., 2017). Edward F. Knipling conceived the concept of SIT, which was first used to control the New World screwworm, *Cochliomyia hominivorax* population, in the 1950s (Bourtzis and Vreysen, 2021).

SIT relies on the production of sterile males to mate with non-sterile wild females, which will, over time, induce a decline in the wild population (Mounika *et al*., 2022). For SIT to be effective, sterile males must be released in an overflooding ratio to the wild population (Carvalho *et al*., 2022). The sterile insect technique (SIT) consists of the mass production, sterilisation, and release of insects into infested areas, where sterile males compete with wild males for mates. Wild females that mate with sterile males will lay fertilised eggs, resulting in no offspring (Osouli *et al*., 2020).

The successful control or eradication of an insect pest requires substantial research and knowledge of the target species, early detection of the pest, and considerable monetary and time commitment (Barnes *et al*., 2015). There are a few success stories where the integration of SIT in AW-IMP has assisted in controlling or eradicating lepidopteran pests (Marec and Vreysen, 2019), such as the Mediterranean fruit fly *Ceratitis capitata* (Wiedemann) (Vreysen *et al*., 2016; Plá *et al*., 2021) and the pink bollworm *Pectinophora gossypiella* (Simmons *et al*., 2020). This method has been successfully used to eradicate or suppress a wide range of insect populations, such as the *C. hominivorax*.

Irradiation technology is the most common method to irradiate insects using gamma or X-rays (Mounika *et al*., 2022). However, due to the genetic makeup of lepidopterans, they are more resistant to radiation technology (Sengupta *et al*., 2022). To achieve the full sterility that is desired for SIT in lepidopterans, very high doses are required; however, such high doses also affect the fitness of the insects, making them incapable of competing with their wild counterparts (Marec and Vreysen, 2019). Nonetheless, lepidopterans are still good candidates for the SIT programmes as inherited sterility is another option; this allows lepidopterans to be irradiated at lower doses to induce partial sterility and fit enough males who, when mated with wild females will pass on radiation-induced deleterious effects to the offsprings rendering the next generation (F1 generation) completely sterile (Saour, 2014; Marec *et al*., 2005). This method has been successfully used to eradicate/control a wide range of insect populations, such as the Mediterranean fruit fly *Ceratitis capitata* (Wiedemann). The F1 generation is male-biased, and because they are produced without irradiation, they are more competitive than the irradiated males, which is a quality suitable for an SIT programme (Vreysen *et al*., 2016;). SIT or inherited sterility has numerous advantages as a pest population control tool, such as species specificity, compatibility with other area-wide control tactics such as the use of GM crops and mating disruption, and is species-specific, ecologically compatible, and non-polluting (Cagnotti *et al*., 2016; Mounika *et al*., 2022).

The sterile insect technique (SIT) is gaining traction as a sustainable and environmentally friendly method for controlling lepidopteran pests. This systematic review examines the role of the sterile insect technique in controlling lepidopteran pests. This is done by investigating publications assessing the application of SIT to suppress or eliminate lepidopteran pests. This review of 24 publications highlights the growing research interest in SIT for Lepidoptera, with a particular focus on (i) geographic distribution of the studies, (ii) target species, (iii) irradiation source (gamma or X-ray), and (iv) when the research has taken place. By summarising information from publications investigating the application of sterile insect techniques for the control of lepidopteran pests, this review can pave the way for future research studies.

## MATERIAL AND METHODS

A systematic review on the application of the sterile insect technique to control lepidopteran pests was conducted following the PRISMA guidelines for a systematic review.

### Search and selection of articles

The following search engines were used to search for peer-reviewed papers published on all continents: Web of Science (http://www.isiwebofknowledge.com), Scopus (https://www.scopus.com), Wiley Online Library (https://onlinelibrary.wiley.com), and JSTOR (https://www.jstor.org). The following search terms were used to find any published material on the application of sterile insect technique for the control of lepidopteran pests: **(“SIT” AND “Lepidoptera” OR “Sterile Insect Technique” AND “Lepidoptera” OR “SIT” AND “Lepidopteran pests” OR “Sterile Insect Technique” AND “Lepidopteran pests”).** The search was limited to papers that were published between 2000 and 2024. The year 2000 was selected because the aim of the paper was to assess research advancements made in the last two decades.

All selected papers were imported as MS Excel files or text files. The initial elimination of the papers was done by reading the titles and abstracts of the selected papers. Once it was determined that the articles were relevant, the full text of the selected articles was further analysed to extract relevant information. The references of the articles selected for full-text reading were also screened to identify studies that might be relevant but did not appear in the initial search.

### Inclusion and exclusion criteria

#### Inclusion criteria

- Original research articles and review papers focusing on SIT as part of management strategies for lepidopteran pests
- Articles published in English
- Articles published between 2000 and 2023

#### Exclusion criteria

- Conference papers, editorial material, book chapters, reports, SIT on other insects
- Articles on other insect orders other than Lepidoptera

### Data quality

The data quality of the selected studies was evaluated based on the following criteria:

(i) a clear description of the insect species studied, (ii) study methods that incorporated SIT as a control strategy for the target pest, and (iii) a study that uses SIT in field or cage trials. (iv) studies that highlight the radiation biology of the insect species. Only studies that met the above criteria were included in the review.

### Data analysis

A narrative synthesis approach was used, and data from the studies were summarised. The results were assessed according to insect species, geographic distribution, and the year the articles were published.

## RESULTS

Figure 1 represents a PRISMA flow chart showing the systematic review process used to select the studies. The search, screening, and inclusion of studies were schematically summarised in the flow chart. The initial search for relevant articles resulted in 2 537 papers from Web of Science (328), Wiley (244), Scopus (225), and JSTOR (1 740) (Table 1). The titles and keywords were screened, and 2 466 articles that did not meet the criteria were excluded. After eliminating duplicates (19 articles), 13 more papers were excluded after reading abstracts. The search criteria yielded 39 articles for full-text reading, and 3 more papers were identified from the references. After full-text reading, 18 papers were excluded, and 24 papers were included in this systematic review.

**Fig 1:**
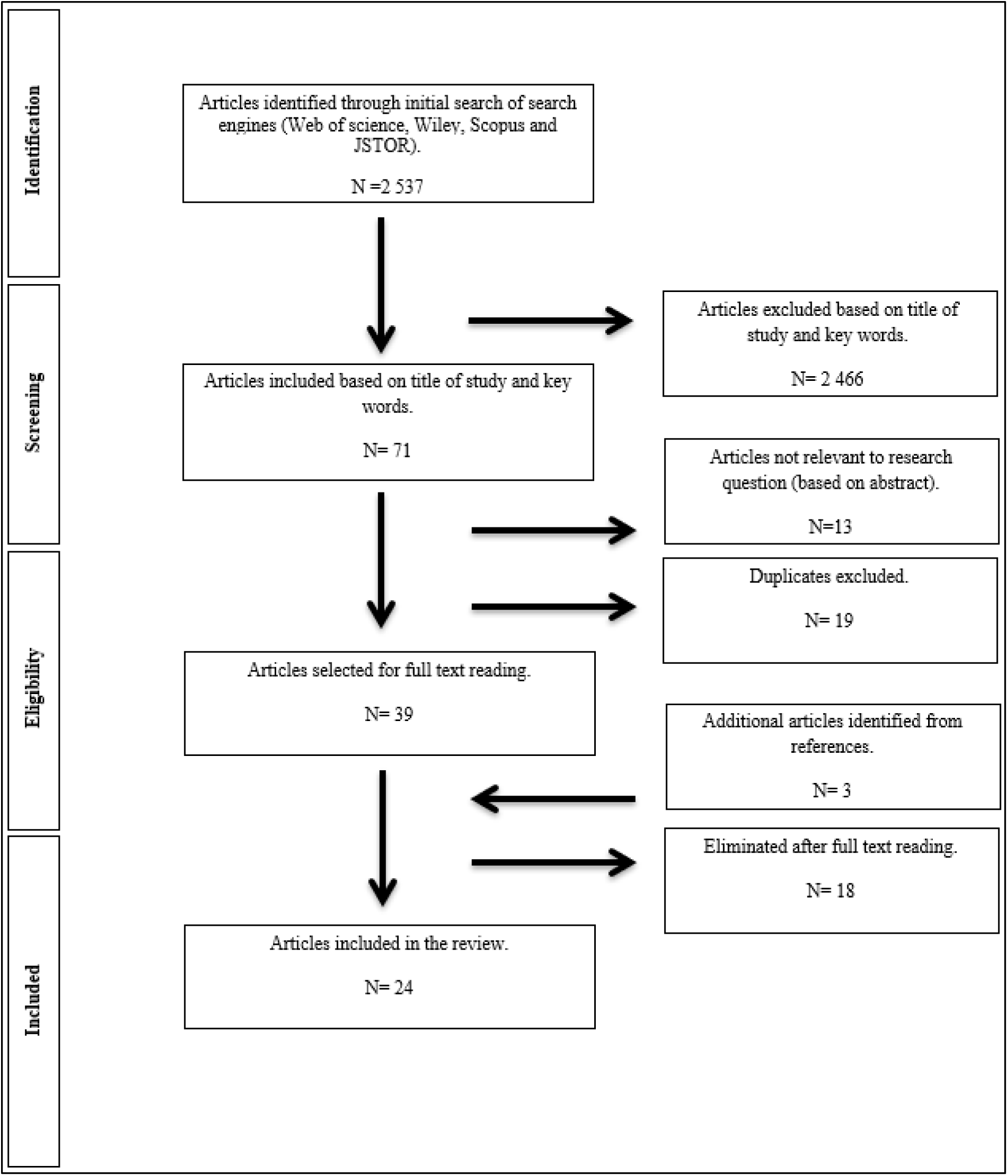
A flow chart showing the systematic review process.

**Table 1:**
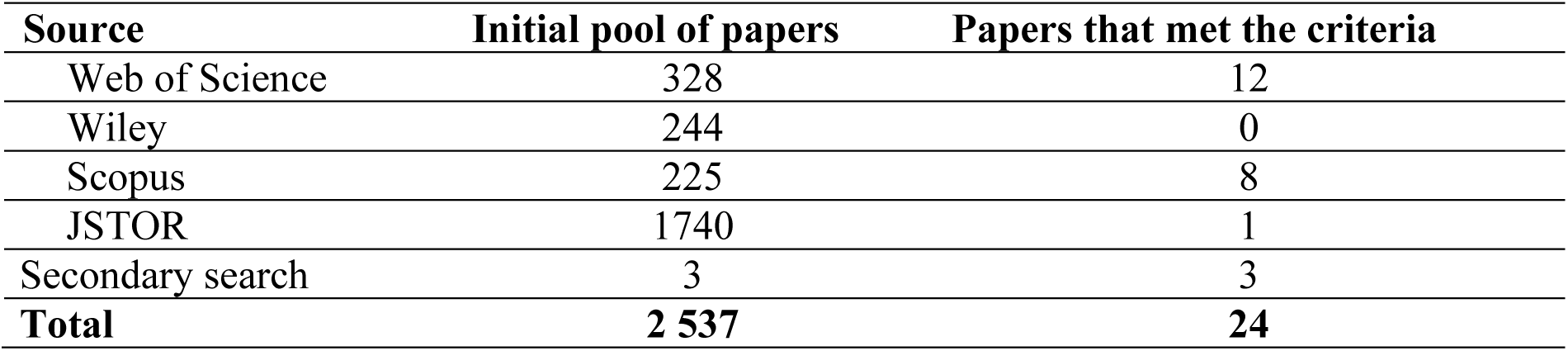
A summary of search and study selection.

### Geographic distribution

Figure 2 represents the geographic distribution of study locations. The selected studies were concentrated in regions with well-established SIT programmes, and research was conducted in developed and developing countries. The highest number of studies were conducted in South Africa (n=6), followed by the United States of America (n=5) and Iran (n=3). These countries have invested significantly in SIT infrastructure and expertise, leading to successful applications against pests like the *Thaumatotibia leucotreta* (false codling moth) and *Pectinophora gossypiella* (pink bollworm). SIT was first developed in the USA and has since been used in six continents (Dyck *et al*., 2021). South Africa is famous for its SIT programme against *T. leucotreta*, and in the USA, *P. gossypiella* was eradicated using SIT (Tabashnik *et al*., 2008).

**Fig 2:**
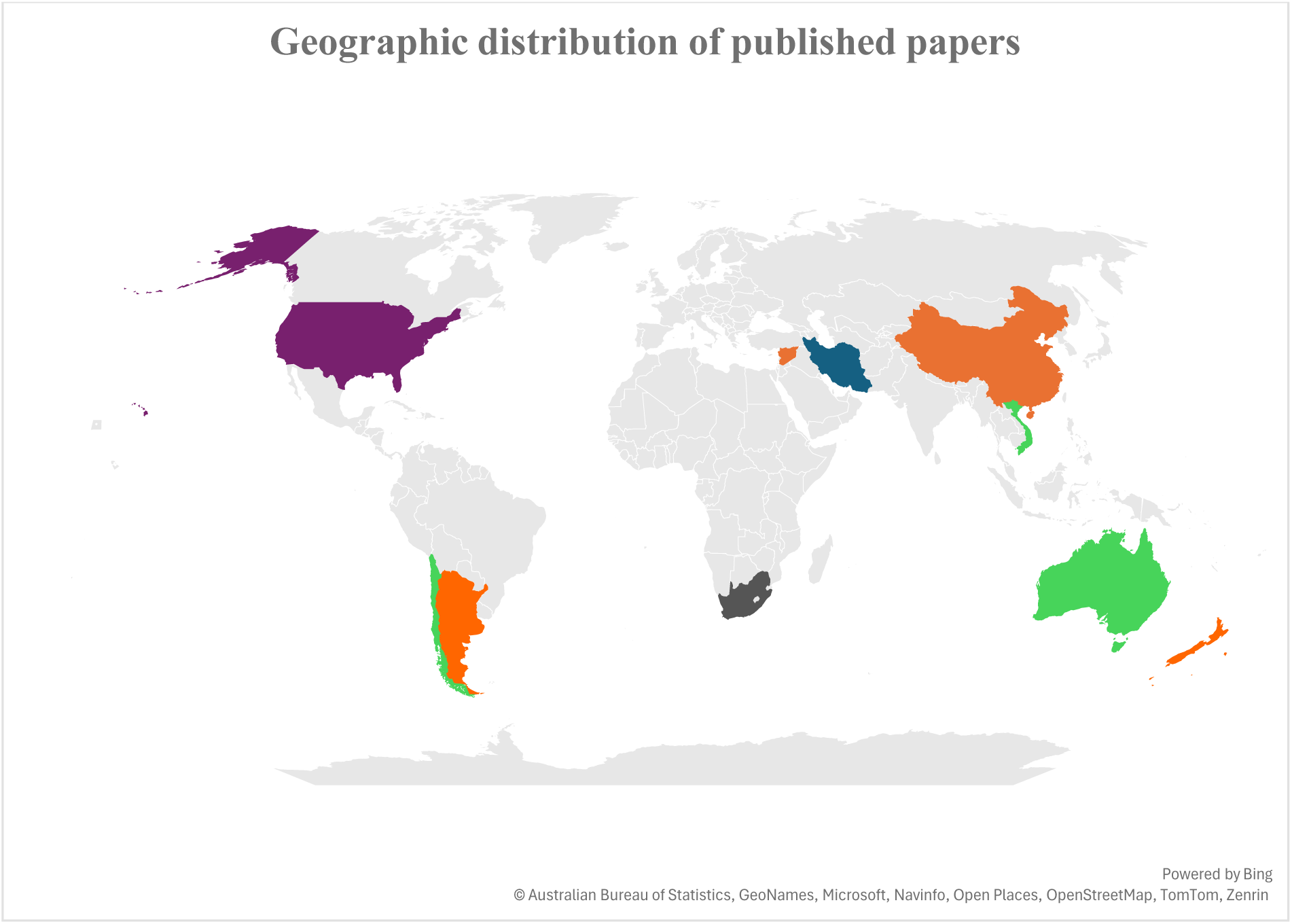
Geographical distribution of the studies assessing the application of sterile insect technique for the control of lepidopteran pests based on 24 primary studies (green=1, orange=2, blue=3, purple=5, Grey=6).

The most studied species, *T. leucotreta*, is native to South Africa and has successfully established itself as a pest. All published papers on this species are from South Africa. A wide range of research on SIT has been conducted in the United States of America on various lepidopteran species such as *Cactoblastis cactorum* (cactus moth) (Carpenter *et al*., 2001, Hight *et al*., 2005;), *Cydia pomonella* (codling moth) (Bloem *et al*., 2001), *Amyelois transitella* (Walker) (navel orangeworm) (Haff *et al*., 2020) and *Epiphyas postvittana* (light brown apple moth) (Jang *et al*., 2012). Fewer studies were conducted in Australia, Chile, and Vietnam (n=1).

### Insect diversity

Table 2 shows the insect diversity, the number of publications, and the irradiation source for each species. Among the 24 publications, 14 species were studied. The families Tortricidae (4) and Pyralidae (4) were the most studied families, followed by the family Gelechiidae (2). The most studied species were the false codling moth *Thaumatotibia leucotreta* (4) and the codling moth *Cydia pomonella* (4). The most studied species in this review was *Thaumatotibia leucotreta* (Lepidoptera: Tortricidae), commonly known as the false codling moth, and all the papers were published in South Africa. The pest is native to sub-Saharan Africa and is found in all citrus production areas in Southern Africa (Coombes *et al*., 2024). It is an important citrus pest in South Africa, and the country is the world’s second-largest exporter of citrus (Roberts & Chisoro, 2021). Because of the quarantine status in markets to which a large portion of the fresh produce is exported, it was essential for South Africa to investigate ways to manage this pest (Coombes *et al*., 2024).

**Table 2:** Insect diversity and the number of papers published for each species.

| Insect species | Number of studies | References |
| --- | --- | --- |
| *Thaumatotibia leucotreta* (Tortricidae) | 4 | Hofmeyr *et al.*, 2016; Hofmeyr *et al.*, 2006; Hofmeyr *et al.*, 2015; Bloem *et al.* 2004 |
| *Cydia pomonella* (Tortricidae) | 4 | Botto & Glaz 2009; Horner *et al.*, 2020; Bloem *et al.*, 2001; Zhang *et al.*, 2022 |
| *Lobesia botrana* (Tortricidae) | 2 | Saour, 2014; Simmons *et al.*, 2021 |
| *Cactoblastis cactorum* (Pyralidae) | 2 | Hight *et al.*, 2005; Carpenter *et al.*, 2001 |
| *Ostrinia nubilalis* (Crambidae) | 2 | Osouli *et al.*, 2020; Osouli *et al.*, 2023 |
| *Epiphyas postvittana* (Tortricidae) | 2 | Soopaya *et al* 2011; Jang *et al.*, 2012 |
| *Eldana saccharina* (Pyralidae) | 1 | Walton & Conlong 2016 |
| *Tuta absoluta* (Gelechiidae) | 1 | Cagnotti *et al.*, 2016 |
| *Plutella xylostella* (Plutellidae) | 1 | Nguyen & Nguyen, 2001 |
| *Amyelois transitella* (Pyralidae) | 1 | Haff *et al* 2020 |
| *Helicoverpa armigera* (Noctuidae) | 1 | Osouli *et al* 2021 |
| *Phyllocnistis citrella* (Gracillariidae) | 1 | Osouli *et al* 2023 |
| *Phthorimaea operculella* (Gelechiidae) | 1 | Makee & Sour 2004 |
| *Ephestia elutella* (Pyralidae) | 1 | Zhao *et al.*, 2022 |

The earliest paper published by Bloem *et al*., 2004 studied the radiation biology of *Cryptophlebia leucotreta* when exposed to different irradiation doses (100-250Gy). It was found that the females were rendered fully sterile at 150-200Gy, while males were partially sterile and could pass on their sterility traits to their offspring. Based on these results, Hofmeyr *et al*. (2006) selected 150 and 200Gy doses to conduct a cage experiment to find the suitable release ratio and examine the suppression capacity of irradiated *C. leucotreta* on a semi-commercial scale. It was noted that a wild-to-sterile release ratio of 10:1 and 5:1 (150Gy and 200Gy, respectively) produced significantly fewer F1 generations. However, the lowest F1 population was produced with 150Gy. From these results, a pilot project was developed for commercialisation at Citrusdal, Western Cape. In 2005, the pilot study continued in commercial citrus orchards in Citrusdal. A mass-rearing facility, which is a subsidiary of Citrus Research International, XSIT (Pty) Ltd, was created in 2007 and still produces sterile *T. leucotreta* for SIT projects (Hofmeyr *et al*., 2015; Hofmeyr *et al*., 2016)

*Cydia pomonella* (Linnaeus, 1758) (Lepidoptera: Tortricidae) was the second most studied species. Papers on this species were published in New Zealand, South Africa, Argentina, and China. In South Africa, *C. pomonella* was first discovered in 1885 and was declared a pest in 1918. South Africa is one of the biggest deciduous fruit producers in the southern hemisphere (Venter *et al*., 2021). According to the systematic review search, the first paper studying this pest was published in South Africa in 2001 (Bloem *et al*., 2001); however, literature reports that the use of SIT on a semi-commercial scale in South Africa for the control of codling moth was officially terminated in 2014 due to a massive budget needed to build proper infrastructures and lack of support from growers.

Honer *et al*. (2020) state that in New Zealand, a six-year SIT programme was initiated from 2014-2019, where an overflooding ratio of 40:1 sterile to wild moths was released in orchards. SIT was used in conjunction with mating disruptions and larvicides, and a population suppression of 90-99% was achieved at the selected release sites. This was done to comply with the export markets’ strict phytosanitary measures and eliminate the risk of larval infestation. Apple growers must heavily control codling moth populations in their orchards to be able to export out of New Zealand. The government of Argentina expressed interest in expanding the use of SIT as part of AW-IPM after it was successfully implemented against fruit flies (Botto & Glaz, 2010). This led to the publication of a study exploring SIT and egg parasitoids to control the codling moth *C. pomonella* in Argentina in 2010. *C. pomonella* is a major threat to the apple industry in China and has become an issue of pome fruit export concern (Wang *et al*., 2019).

This pest is native to central Asia but has spread throughout China (Chen *et al*., 2020). This has motivated research into the study of its radiation biology to delay its further spread into the main production areas.

Zhang *et al*. (2022) studied the effect of different irradiation doses on the sterility of *C. pomonella* F1 generation. It was deduced that an irradiation dose of 366Gy was suitable when an X irradiator was used (Zhang *et al*., 2022). Earlier work on *C. pomonella* was done by Bloem and Bloem (1996), which led to the actual implementation of the SIT programme, the Okanagan-Kootenay Sterile Insect Release Programme for codling moths (Bloem *et al*., 2001). The facility has been operating since 1992, producing approximately 2 million sterile moths daily. It also ships sterile insects to other countries. In the past 20 years, it has achieved a 94% population reduction (Nelson *et al*., 2021).

In this case, the initial source of irradiation was Cobalt60 (60Co-γ); however, due to health risks and problems associated with acquiring a gamma irradiator, an alternative irradiation source had to be identified. (Odendaal *et al*., 2015). The European grapevine moth, *Lobesia botrana* (Denis and Schiffermuller) (Lepidoptera: Tortricidae), is a polyphagous insect that develops on more than 200 plant species from various families; nonetheless, it remains a major pest in vineyards. This resulted in the publication of two papers on radiation biology and the potential usefulness of SIT in controlling *L. botrana*. Sour (2014) outlines the effect of irradiation on *L. botrana* and has provided a starting point for developing an SIT programme on this pest species. According to this study, 150Gy was an appropriate dose for male European grapevine moths to pass their deleterious effects to the F1 generation. In 2015, an SIT programme was developed in Chile to evaluate the effectiveness of SIT in the field when sterile moths are released in Urban areas adjacent to agricultural farms and to confirm the 150Gy dosage for inherited sterility (Simmons *et al*., 2021). Results from this study were inconclusive about the effectiveness of SIT; 150Gy was confirmed to be an adequate dose for implementation of Inherited sterility. However, it was deduced that this area had a large population of wild moths that needed to reach the sterile-to-wild moth ratios needed to achieve effective control.

Before the cactus moth *Cactoblastis cactorum* (Berg) (Lepidoptera: Pyralidae) was declared a pest in the United States of America; it was used as a biological control agent against weeds as it played a role in obliterating invasive *Opuntia* in Australia in the 1930s. However, when it was introduced in the United States of America, concerns were raised about many species of native *Opuntia* in southern North America (Height *et al*.,2005). Carpenter *et al*. (2001) published the first article studying the irradiation of *C. cactorum* since its introduction in the United States and Mexico. This was encouraged by the observed F1 sterility proven in numerous economically important Lepidoptera (Bloem & Carpenter, 2001).

The study suggested that *C. cactorum* is a good SIT candidate; however, a suitable dose needed to induce full sterility in the F1 generation was not confirmed but ranged from 100-200Gy (Carpenter *et al*., 2001). In a study conducted by Hight *et al*., 2005, the effect of releasing different ratios of partially sterilised males and fully sterilised females in field cages containing potted host plants and a cohort of untreated *C. cactorum* males and females was evaluated. Both males and females were treated to an irradiation dose of 200Gy and released at a ratio of 5:1. This was enough to decrease the population of wild in field cages irrespective of whether only partially sterile males were released alone or together with fully sterile females (Hight *et al*., 2005).

These results provided valuable evidence that an SIT/F1 sterility program can be initiated toward controlling *C. cactorum*. Osouli *et al*., 2020 studied the effect of different doses of gamma radiation on the biological and reproductive parameters of *Ostrinia nubilalis* Hubner (European corn borer) (Lepidoptera: Crambidae), a key pest of grain corn and sweet corn in Iran. Results showed that an irradiation dose higher than 300 or 350 Gy would be adequate for inducing proper sterility in the female and male of *O. nubilalis,* respectively. The results did, however, show that it would be more beneficial to release male parents that were exposed to sub-sterilizing doses as pupae as the progeny from which were more sterile than the parents. The presented results set the ground for developing an SIT programme as a component of IPM against *O. nubilalis* in 2023. Osouli *et al*. (2023) published a study on the effect of releasing sterile moths in experimental cages and observed that releasing sterile *O. nubilalis* at a sterile-to-wild ratio of 7:1 decreased the mean percentage of stem infestation on maize plants by up to 14.2%.

The light brown apple moth *Epiphyas postvittana* Walker (Lepidoptera: Tortricidae) is native to Southeastern Australia, but it has gradually established itself in other parts of Australia (Thrimawithana *et al*., 2022). It has also established itself as a pest in New Zealand. In both New Zealand and Australia, it is an important economic pest of grapes, citrus, apples, and pears (Murray, 2014). Because of its economic importance in these two countries, coordinated radiation biology studies were undertaken on two geographically isolated light brown apple moth populations from Australia and New Zealand to initiate a full-sterility, or F1 inherited sterility program against light brown apple moth (Soopaya *et al*., 2011). The moths from these two areas were exposed to the same irradiation and subjected to the same laboratory tests. The results showed that in both populations, when *E. postvittana* pupae were exposed to an irradiation dose of 250 Gy, 95% and 90% sterility was induced in females and males, respectively. This suggests that an SIT or F1 sterility program can be applied to control *E. postvittana*; however, further research needs to test the fitness and field performance of irradiated moths as well as the research on modelling the overflooding ratios needed for effective control. In 2012, Jang *et al*. further expanded on the research on *E. postvittana* by conducting a study on F1 hybrid sterility to assess if higher irradiation doses needed for full parental generation sterility could reduce to increase fitness. The results did, however, show that a dose of 250 or 300 Gy was adequate for the application of SIT, whereby few or no progeny was found in irradiated adult moths and that further research on F1 hybrid sterility is needed (Jang *et al*., 2012). These results corroborated the findings reported by Soopaya *et al*., 2011.

Sugarcane stalk borer *Eldana saccharina* Walker (Lepidoptera: Pyralidae) is a native pest to Africa, occurring on many gramineous crops (Walton and Conlong, 2016b). It has been a major pest in the South African sugar industry for many years and causes enormous economic injury (Mulcahy *et al*., 2023). It has been reared for many years in the South African Sugarcane industry, and Walton and Conlong (2016a) studied its irradiation biology to establish an SIT programme against *Eldana Saccharina*. It was found that, like most lepidopterans, male *Eldana Saccharina* can be irradiated at a sub-sterilizing irradiation (200Gy) and pass on the sterility to the F1 generation (Walton & Conlong, 2016a).

*Plutella xylostella* (L.) (Lepidoptera: Plutellidae), also known as the diamondback moth (DBM), is a serious pest of cruciferous crops throughout the world. Earlier studies by Omar & Mansor (1993), Sutrisno *et al*. (1993), and Sutrisno & Hoedaya (1993) report that the radiation doses between 150 Gy and 200 Gy were suitable for inducing inherited sterility in DBM. Based on the findings of these studies, a field-based study was conducted to explore the potential suppression of DBM populations using the Fl sterility technique in 2001 (Nguyen & Nguyen, 2001). Parental insects were treated with a dose of 200 Gy and outcrossed with untreated counterparts. The F1 and F2 resulting from the parental crosses were also outcrossed with wild insects, and the effect of different release ratios of the parental, F1, and F2 diamondback moths was observed in cages containing planted cabbage. The sterility of treated males was 62.1%, Fl males was 94.8%, and F1 females was 91.1%. In the F2 generation, sterility ranged from 61.8% to 64.8%. The paper further reports that releasing the parasitoid *Cotesia plutellae* in combination with inherited sterility may be feasible for managing early-season populations of DBM. *Tuta absoluta* (Meyrick) (Lepidoptera: Gelechiidae) is a tomato pest that is native to South America. It is now considered a key agricultural threat to European and North African tomato production as it has spread throughout the world since its first discovery as a prolific pest in Spain in 2006 (Cagnotti *et al*., 2016). In a field cage experiment set up in Argentina, releasing sterile moths, it was confirmed that releasing sterile *T. absoluta* at a ratio of 15:1 caused a population decline over time (Cagnotti *et al*., 2016).

*Amyelois transitella* (Walker) Navel Orangeworm (Lepidoptera: Pyralidae) causes damage to tree nuts, including mostly almonds, walnuts, and pistachios, and has been an expensive pest to California’s agricultural department. SIT has not yet been used for control of navel orangeworm in California orchards; therefore, Haff *et al*. (2020) conducted a study to test the potential of using X-ray to irradiate *A. transitella* pupae and larva for SIT purposes as previous studies using gamma had reported that they were not suitable candidates based on high mortality rates (Husseiny & Madsen, 1964). Although low mortality was observed in this study, the use of the absence of offspring as the measure of sterility allowed a simpler and less labour-intensive means to determine the potential suitability of Navel Orangeworm larvae and pupae for SIT under x-ray irradiation as opposed to gamma, at the end the study was deemed inconclusive and warrants further research (Haff *et al*., 2020).

Old-world bollworm *Helicoverpa armigera* Hübner (Lepidoptera: Noctuidae) has established itself as a pest in Iran. It has been reported on over 180 cultivated hosts and wild species and is a major pest of cotton, tomato, maize, chickpea, and pistachio (Atashi *et al*., 2021). Pistachio is among the most important export products of Iran, and due to the increasing infestation of old-world bollworms, research was initiated to study the effects of sterilizing and sub-sterilizing doses of gamma radiation on *H. armigera* (Osouli *et al*., 2020). Treating *H. armigera* with increasing radiation doses reduced the fertility of both male and female moths, and at 350Gy, no offspring were produced. This dose can be used for irradiation and release of parental moth and provides a foundation to promote the development of the SIT programme.

Citrus leafminer (CLM), *Phyllocnistis citrella* Stainton (Lepidoptera: Gracillariidae), is a key pest of young citrus, a major problem in citrus nurseries. Citrus leafminer is native to Asia but has colonised many countries, including Iran. The field performance of sub-sterilised citrus leafminer was evaluated by Osouli *et al*., 2023. Different release ratios of sterilised citrus leafminers were released in cages to assess plant infestation and determine the suitable release ratio for an SIT against this moth. In this study, an irradiation dose of 250Gy resulted in full sterility for both sexes of CLM, and a release ratio of 20:1 caused a 46.1% reduction in larval infestation (Osouli *et al*., 2023).

*Phthorimaea operculella* Zeller (Lepidoptera: Gelechiidae) is an important insect pest of cultivated potatoes worldwide. Little information was available on this species, and in 2004, Makee and Sour studied the radiation biology of the potato tuberworm. Complete sterility in females was obtained at 200 Gy, whereas a dose above 400 Gy was required in the males; just like in many other lepidopterans, the radiosensitivity of P. operculella females was higher than that of males. A noticeable reduction in population was observed when a 1:1:10 ratio of irradiated males was used, and satisfactory results were reached when a high ratio of partially sterile males was used (Makee and Sour, 2004).

The tobacco moth, *Ephestia elutella* (Hübner) (Lepidoptera: Pyralidae), is responsible for approximately 1% of the annual loss of stored tobacco. It is a polyphagous pest infesting stored products such as tobacco, nuts, dried fruits, and cereals. A lot of research done on the management of this species relies on the use of insecticides. However, new control strategies are needed, so this study (Zhao *et al*., 2022) investigated the optimal irradiation dose of X-rays to control *E. elutella*. A dose of 200Gy was suggested as the appropriate dose to induce sterility in males; as a result, X-ray irradiation at 250 Gy for the control of *E. elutella* infestation should be further tested, and future research should determine the feasibility and effectiveness of X-ray (Zhao *et al*., 2022).

### Irradiation source

Gamma (n=20) was the most used irradiation source to sterile the studied species, while only four studies were conducted using the X-ray source (Table 3). The *C. pomonella* was the only species studied using the gamma and the X-ray sources. Although gamma radiation is the most commonly used method for sterilising insects, the high cost and safety concerns associated with gamma irradiators have led to increasing interest in alternative irradiation sources, such as X-rays (Yamada *et al*., 2023).

**Table 3:**
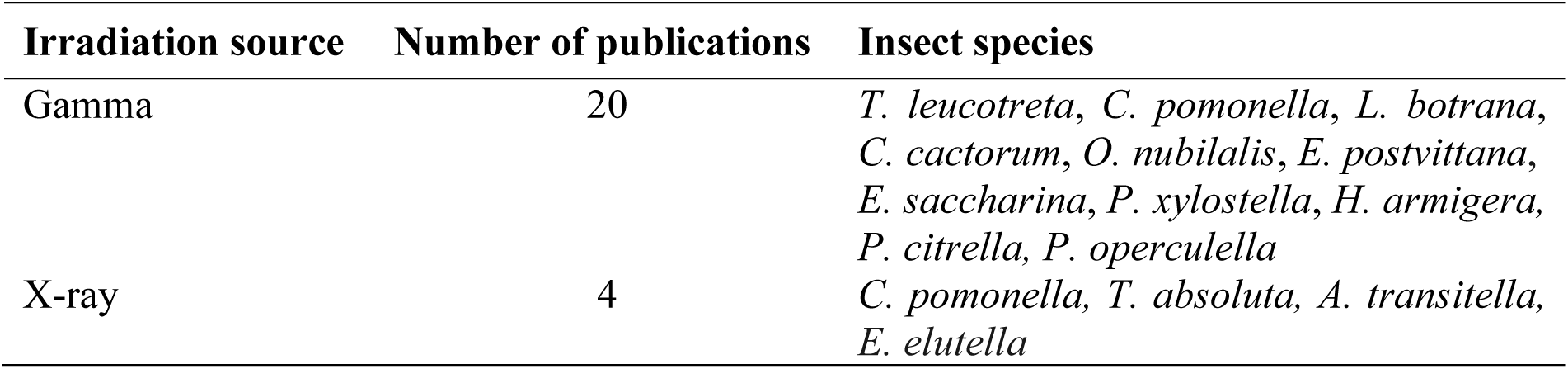
Irradiation source used to sterilise the insect species.

| Irradiation source | Number of publications | Insect species |
| --- | --- | --- |
| Gamma | 20 | *T. leucotreta*, *C. pomonella*, *L. botrana*, *C. cactorum*, *O. nubilalis*, *E. postvittana*, *E. saccharina*, *P. xylostella*, *H. armigera*, *P. citrella*, *P. operculella* |
| X-ray | 4 | *C. pomonella*, *T. absoluta*, *A. transitella*, *E. elutella* |

### Sterility technique

Two common sterility techniques were used in the selected papers (Table 4). These were the traditional sterile insect technique, where insects are mass-reared, irradiated, and released into the field, as well as the inherited sterility (F1 sterility). F1 sterility is similar to the sterile insect technique as it also involves the mass rearing of the target species; however, in this case, parents are partially irradiated and allowed to mate and produce a completely sterile F1 generation that will be released in the field. This technique offers a promising alternative, and technique allows for the use of lower radiation doses, potentially improving the competitiveness of released insects.

**Table 4:**
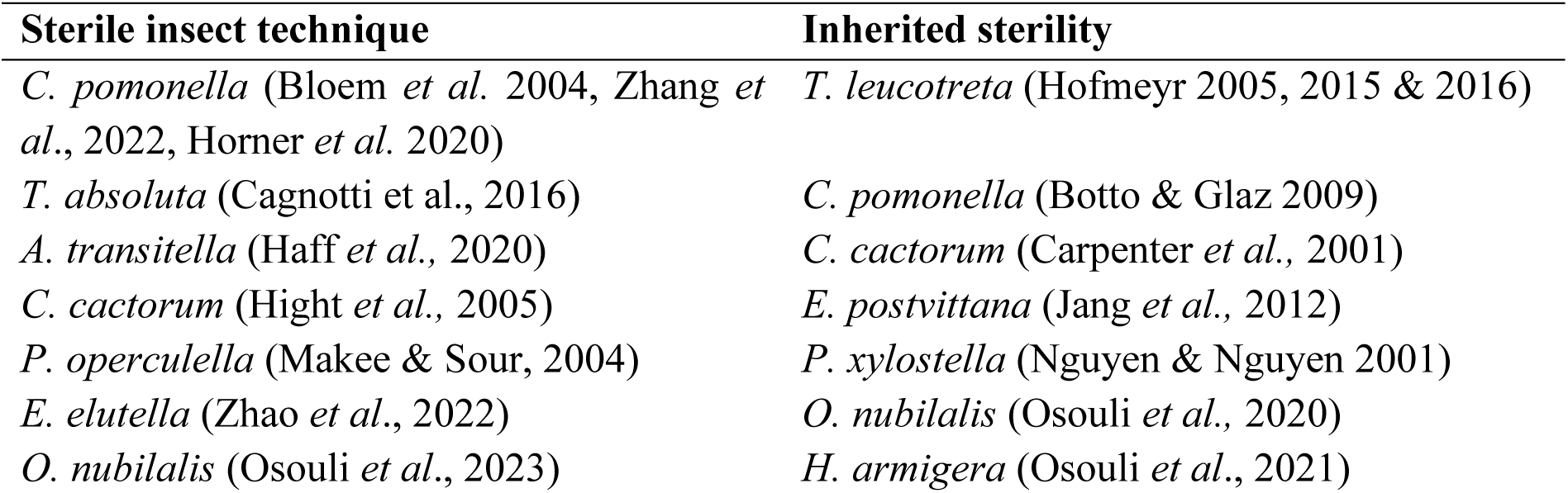
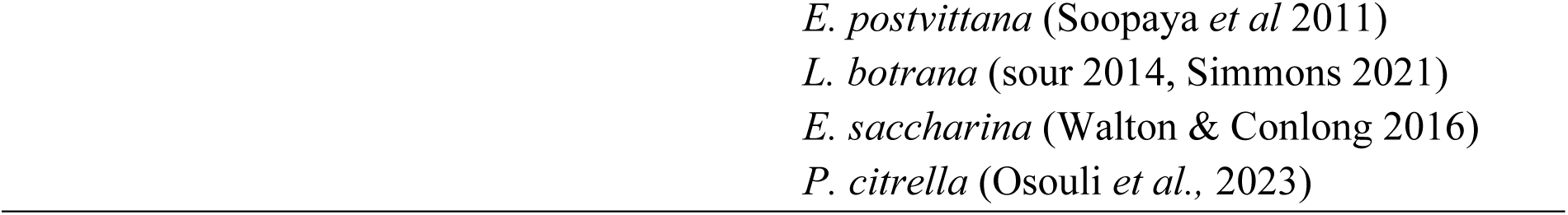
Sterility technique used to treat the insects.

| Sterile insect technique | Inherited sterility |
| --- | --- |
| *C. pomonella* (Bloem *et al.* 2004, Zhang *et al.*, 2022, Horner *et al.* 2020) | *T. leucotreta* (Hofmeyr 2005, 2015 & 2016) |
| *T. absoluta* (Cagnotti *et al.*, 2016) | *C. pomonella* (Botto & Glaz 2009) |
| *A. transitella* (Haff *et al.*, 2020) | *C. cactorum* (Carpenter *et al.*, 2001) |
| *C. cactorum* (Hight *et al.*, 2005) | *E. postvittana* (Jang *et al.*, 2012) |
| *P. operculella* (Makee & Sour, 2004) | *P. xylostella* (Nguyen & Nguyen 2001) |
| *E. elutella* (Zhao *et al.*, 2022) | *O. nubilalis* (Osouli *et al.*, 2020) |
| *O. nubilalis* (Osouli *et al.*, 2023) | *H. armigera* (Osouli *et al.*, 2021) |

### Effectiveness of SIT

Only recently has SIT research made its way out of the laboratory and into the field. The search results report that the first paper publishing results from SIT research conducted in the cages/field was in 2001. In the first paper that was published in 2001, the influence of introducing a population of sterile *Plutella xylostella* in a cage with planted cabbage at a different overflooding ratio of sterile to wild *P. xylostella* was studied. Results showed that the population of *P. xylostella* was significantly reduced when sterile males were released into the cage in a 5:1 sterile to wild overflooding ratio. Compared to the control, there was a significant reduction in the number of F2 adults emerging in the field cages receiving sterile moths. A 68% reduction in the F2 generation was observed for field cages receiving irradiated moths (Nguyen & Nguyen, 2001).

In a study conducted by Cagnotti *et al*., 2016 a release ratio of over-flooding ratio of 10:1 sterile to wild *Tuta absoluta* (Gelechiidae) caused a decline in larval production in treatment cages containing potted tomato plants as compared with the untreated control group. Furthermore, it was noted that increasing the treated: untreated moth ratio to 15:1 resulted in significantly fewer small and large larvae than the untreated control cages across the 3 generations studied. The sterilised males were as competitive as the wild males and were also successful in producing sterile F1 progeny that reduced the wild population growth over time (Cagnotti *et al*., 2016).

Similar results were observed when an over-flooding ratio as low as 5:1 (treated: fertile) of *Cactoblastis cactorum* (Pyralidae) was released in experimental cages. A ratio of 10:1 was even more effective in reducing the percentage of egg hatch and the number of larvae produced in the next generation (Hight *et al*., 2005). A paper by Osouli *et al*., 2023, shows the effect of releasing sterile *Phyllocnistis citrella* and *Ostrinia nubilalis* in field cages. There was a 46.1% reduction in larval infestation of citrus leaves in cages receiving sterile *P. citrella* and a 75,5% reduction in larval infestation of maize plants in cages receiving sterile *Ostrinia nubilalis*. SIT was successful in reducing the population of wild *P. citrella* and *O. nubilalis* in field cages.

In an unconventional study by Simmons *et al*., 2023, Sterile insects were released in an urban area of Chile next to a mixed agricultural crop region, which included grapes, berries, tree fruits, and mixed field crops to control populations of *Lobesia botrana*. Evaluating the effectiveness of SIT in this study was difficult, as the results were the only data available from the recapture of *L. botrana* in traps strategically placed in the urban area. The research focused on dispersal studies of irradiated moths rather than the effectiveness of using SIT to control the wild population; however, the results showed a lower number of wild moths in the release field than in the control field. The results were inadequate to conclude the effectiveness of releasing SIT (Simmons *et al*., 2023).

A research project on season-long releases of partially sterile *Cydia pomonella* conducted in an apple orchard on Lake Osoyoos in North America was also one of the first papers to be published on the use of SIT in the field. However, the results of this project were inconclusive. The season-long releases of partially sterile *Cydia pomonella* (Tortricidae) males showed no significant difference in fruit damage between the treated plots and the control at harvest. It was deduced that this could have been due to the migration of the released sterile males from the treated plots into the control. There was evidence of sterile males in the control plot while monitoring the fields (Bloem *et al*., 2001).

This movement of the sterile males influenced the results inside control plots at two different sites and might render the results inconclusive. However, at one of the selected release sites, fruit damage at harvest decreased from 2–1%, which indicates that the release of partially sterile moths can be a very effective control tactic. Bloem *et al*., 2001) suggested that if tree plantings are contiguous, a distance of at least 500 m is needed to eliminate the possibility of insects entering the control plot.

*Cydia pomonella* was, however, suppressed in a six-year-long SIT project conducted in New Zealand (Honer *et al*., 2020), where sterile insects were released at a 40:1 sterile-to-wild ratio in orchards. It is worth noting that the release of sterile males in this project was used in combination with mating disruption and the application of insecticides, and thus, it cannot be concluded that SIT was effective. However, SIT is often most effective when integrated with other area-wide integrated pest management (AW-IPM) strategies, such as mating disruption and the use of biological control agents. This integrated approach provides a more comprehensive and sustainable pest management solution.

Research on developing an SIT programme against *Thaumatotibia leucotreta* was conducted, leading to the actual implantation of an SIT project that was fully commercialised (Hofmeyr *et al*., 2016). Initial research showed that the overall mean crop loss to *T. leucotreta* infestation during the experimental period was 0.1 and 2.1 infested fruit per tree per week in the SIT and control sites, respectively, representing an infestation reduction of 95.2% using SIT (Hofmeyr *et al*., 2015).

Currently, SIT/IS programmes rely on the release of both males and females, but evidence has shown the economic benefits of male-only releases by decreasing production costs and increasing the efficiency of males (Marec *et al*., 2005). This was well documented in the SIT programme against the mediterranean fruit fly where the manipulation of a genetic sexing strain allowed for the mass rearing and releasing of males only (Rendón *et al*., 2004). This gene is not expressed in many lepidopteran species; however, a pure genetic sexing system was developed for two lepidopteran species: *Bombyx mori* Lepidoptera: Bombycidae and *Ephestia kuehniella* Lepidoptera: Pyralidae (Strunnikov, 1975; Ohnuma, 2005 & Marec 1991).

This system is based on the construction of a balanced lethal strain generating trans-heterozygous males for two sex-linked recessive lethal mutations in such a way that when males are mated to wild females, the resulting offspring are exclusively males only. In recent years, new opportunities have emerged for genetic sexing along with advancements in genetic technologies, especially in transgenesis and gene editing (Marec & Vreysen, 2019). It has been proposed that a genetic sexing strain can be constructed using a transgenic approach so that females can carry a transgene with a dominant conditional lethal mutation in their W chromosomes that would be expressed in embryogenesis. The eggs would then produce non-transgenic male-only progeny that could be released (Marec *et al*., 2005). While recent advancements in genetic techniques have opened new avenues for SIT, there is limited research on how these innovations can be systematically integrated into existing SIT frameworks to improve the competitiveness and viability of sterile insects; as a result, further research into this field is warranted.

Table 5 shows the effectiveness of SIT when applied to semi-field and field conditions. SIT was proven to be effective in controlling populations of target pests in field conditions by eight of these studies, while the results from three of the studies were inconclusive because SIT was used in combination with other AW-IPM methods, such as insecticides and parasitoids.

**Table 5:** The effectiveness of the sterile insect technique when applied in cage or field trials.

| Effective | Inconclusive |
| --- | --- |
| *T. absoluta* (Cagnotti *et al.*, 2016) | *C. pomonella* (Bloem *et al.*, 2001; Honer *et al.*, 2020) |
| *P. xylostella* (Nguyen & Nguyen, 2001) | *L. botrana* (Simmons *et al.*, 2021) |
| *C. cactorum* (Hight *et al.*, 2005) |  |
| *P. citrella* (Osouli *et al.*, 2023) |  |
| *O. nubilalis* (Osouli *et al.*, 2023) |  |
| *T. leucotreta* (Hofmeyr, 2005, 2025 & 2016) |  |

### Trends in the year of publication

The number of publications on SIT for Lepidoptera has increased steadily over the past two decades. This indicates a growing recognition of SIT’s potential and the need for continued research to refine and expand its application. According to the systematic review, the first papers addressing the application of the sterile insect technique to control lepidopterans in the last two decades were published in 2001, with no papers published from 2002-2003 (Figure 3). Since then, there has been an increase in interest in this research field as at least one paper has been published every other year after 2004. Although SIT was initially used to eradicate a target pest, it is increasingly sought as a population suppression method with growing interest from commercial insect producers (Kapranas *et al*., 2022). Because the interest in SIT as a research field is gaining traction, an alternative irradiation source other than the universally and commonly used gamma irradiator had to be found (Mudavanhu *et al*., 2017). This is because the gamma irradiator is expensive, very difficult to acquire (constraints of transportation, import legislation), and is associated with many health and safety concerns to humans, thus limiting the use of gamma irradiators for large quantity producing SIT programmes (Ndo *et al*., 2014; Yamada *et al*., 2023; Kaboré *et al*., 2023). X-rays in the last decade have gained traction, especially from smaller or start-up SIT projects, due to simplified procurement procedures and the absence of safety regulations in the importing country (Yamada *et al*., 2023). Figure 3 below reviews the number of selected studies published on the application of this technique over the past two decades. Most of the studies were conducted during the last decade.

**Fig 3:**
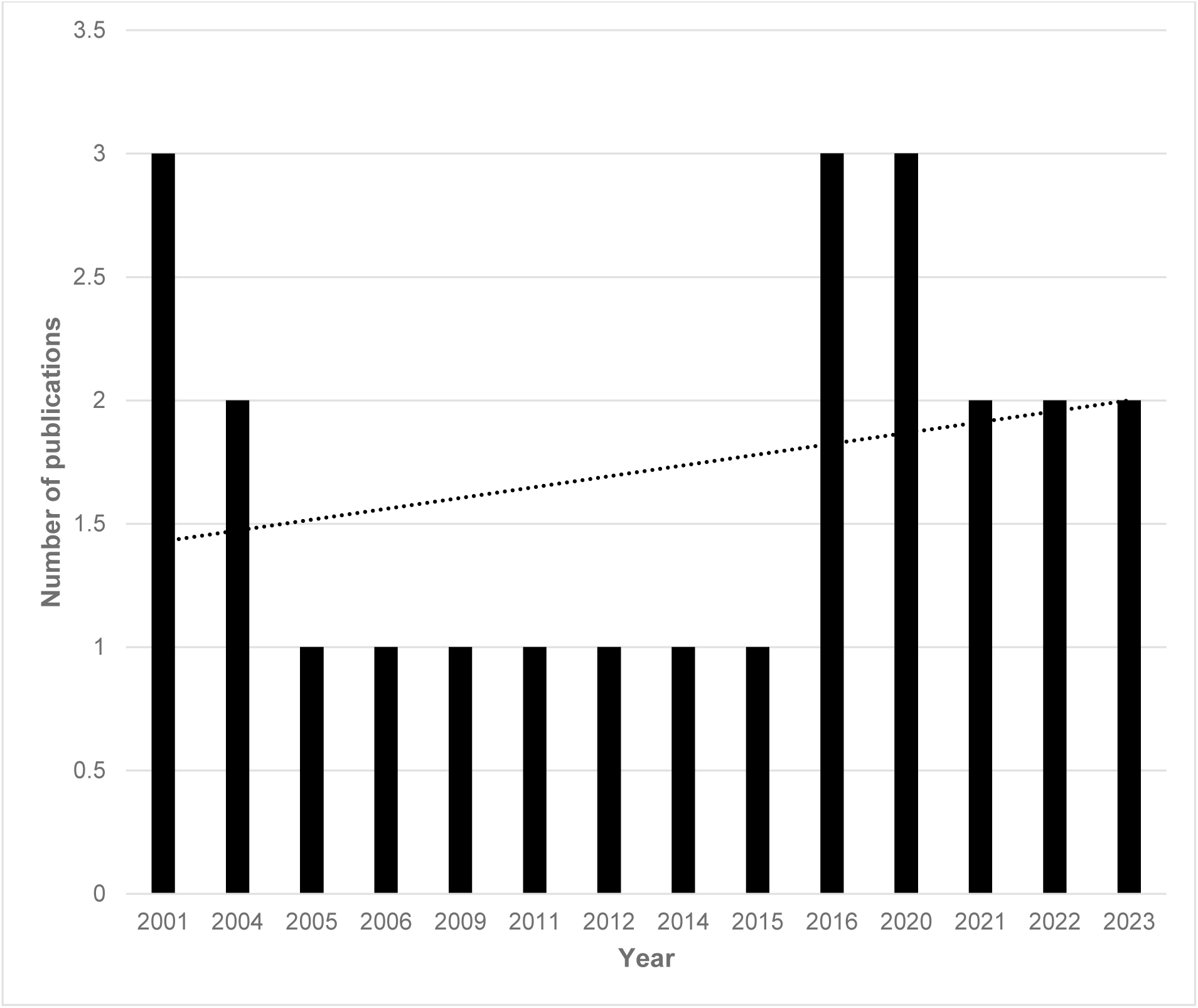
Number of studies assessing the application of sterile insect technique for the control of lepidopteran pests published between 2001-2023.

### Methodological approach

Figure 4 represents the study approaches used for the 24 selected publications. The predominant type of the studies was performed in laboratory settings (58%), while 25% were conducted in semi-field (cage trials) conditions and 17% under field conditions.

**Fig 4:**
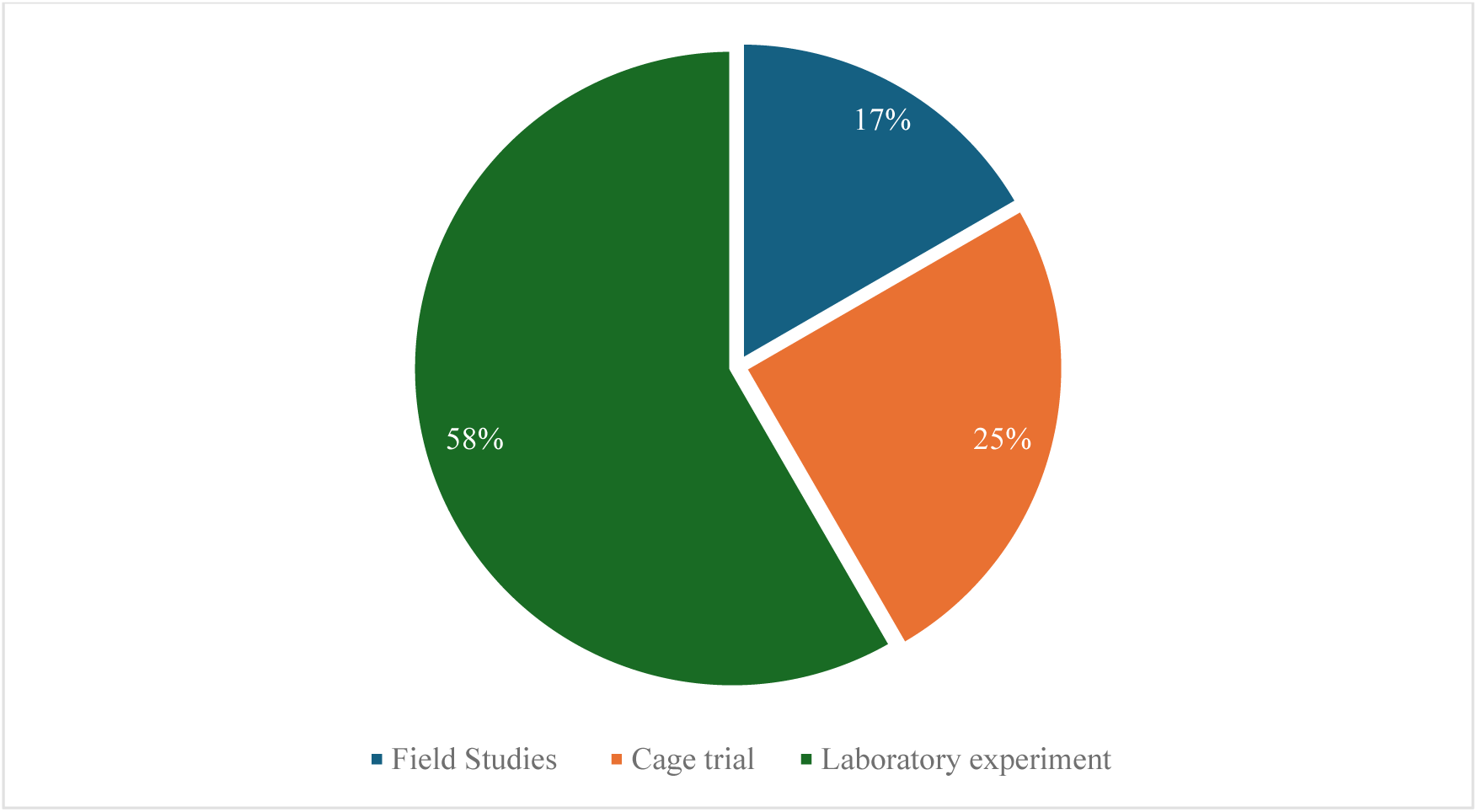
Types of studies/experiments in published selected articles.

From the 24 articles selected for the systematic review, 15 of the papers report on the lab experiments. This included studying irradiation biology and the effect of irradiation on the fertility and fecundity of the insects. The papers focusing on lab experiments were published between the years 2001 and 2023. For a sterile insect technique project to be implemented in a field study or cage trial, an appropriate dose for sterilisation must be established first, and the studied insect must remain fit and able to mate after receiving irradiation.

As a result, most papers published evaluated the suitability or potential of lepidopterans for sterile insect technique and were conducted in a laboratory setting. After a species has proven to be a good SIT candidate, its fitness and ability to compete in the field must be tested before it can be released. This is done by designing cage trails that offer conditions similar to those in the field in which said species will be released. From the search results obtained, 5 articles evaluated the performance of the studied species in semi-field conditions.

Cage trials are also important in determining the release ratio needed for population suppression in the field. However, it is impossible to know for certain the wild population in the field. Research from cage trials provides baseline research for implementing field releases. The papers focusing on cage trials were published between 2005 and 2023. Although research on SIT started in the late 1900’s, it was only recently that cage and field studies were established.

SIT field releases were implemented on a commercial scale in 2001. Since the start of SIT research in the late 1900s, a few SIT programmes have been fully implemented in British Columbia (the Okanagan-Kootenay Sterile Insect Release (OKSIR)), Chile, New Zealand, and South Africa. Five papers from this systematic review report on using the Sterile Insect technique as part of area-wide integrated pest management programmes to control various lepidopteran species both on a small and commercial scale.

## CONCLUSION

The sterile insect technique (SIT), which comprises overflooding releases of sterile insects to suppress or eradicate the target pest, has been reported to be an effective area-wide integrated pest management method. Although the technique relies on continuously releasing sterilised insects, it is an environmentally friendly pest management technology. The initial search of the databases resulted in 2 537 papers, with many papers being either reports or conference proceedings; thus, they were eliminated from the study. The selected papers reported on the application of SIT in the field, cage trials, or laboratories. Twenty-four papers were included in this review, and 14 lepidopteran species were distributed in 10 countries.

The application of sterile insect technique for the control of lepidopteran pests has been well studied over the past two decades. Extensive research has been conducted on the radiation biology of various lepidopteran species. The false codling moth (*Thaumatotibia leucotreta*) and codling moth (Cydia pomonella) are the most studied species. These pests pose significant economic threats to fruit production, driving extensive research efforts to develop effective SIT control strategies. The irradiation dose required to irradiate the insect’s species ranged from 100Gy-400Gy across all species.

The dose 200Gy was commonly used as a desired irradiation dose. The lower radiation dose used to partially sterilise males results in more active insects, disperses greater distances, and is generally more competitive (Bloem *et al*., 2001). It is worth noting that the use of gamma as an irradiation source is now phased out, and X-ray is becoming popular. Successful examples include control of the diamondback moth, tomato leafminer, and cactus moth (Hofmeyr *et al*., 2016; Hofmeyr *et al*., 2015; Honer *et al*., 2020; Bloem *et al*., 2001; Simmons *et al*., 2021).

However, some studies report inconclusive results, particularly when SIT is integrated with other pest management techniques. There is a lack of adequate scientific papers on the application of SIT in the field that needs to be further explored. While laboratory studies dominate the literature, there is a growing trend towards field and cage trials. This shift reflects the increasing emphasis on evaluating the practical application and effectiveness of SIT in real-world settings. Therefore, it is important to improve understanding of the dispersal and mating behaviour of irradiated insects in different environments. This information is crucial for optimising release strategies and ensuring effective population suppression. It is also important to develop robust monitoring and evaluation tools to accurately assess the impact of SIT on target pest populations and non-target organisms.

The sources consistently emphasise the importance of continued research to optimise SIT for different lepidopteran species and to develop cost-effective and accessible irradiation technologies. Further research is also needed to enhance public awareness and engagement to promote the adoption and acceptance of SIT as a safe and environmentally friendly pest control method. By addressing these research priorities, SIT can become an even more powerful tool for the sustainable and effective management of lepidopteran pests worldwide.

